# UHRF1 Overexpression Generates Distinct Senescent States with Different Tp53 Dependencies

**DOI:** 10.1101/2025.11.26.690503

**Authors:** Elena Magnani, Charlene Chen, Filippo Macchi, Bhavani P Madakashira, Tijana Randic, Ian McBain, Kirsten C. Sadler

**Affiliations:** Program in Biology, NYU Abu Dhabi, PO Box 129188, Abu Dhabi, United Arab Emirates; Center for Genomics and Systems Biology, NYU Abu Dhabi, PO Box 129188, Abu Dhabi, United Arab Emirates

**Keywords:** Senescence/DNA methylation/tp53/zebrafish/preneoplastic/liver cancer

## Abstract

Senescence is a pleiotropic phenotype that alternatively suppresses or promotes cancer. The tumor suppressive roles are linked to clearance of damaged cells by the immune system, while cells with tumor promoting functions resist apoptosis, evade immune clearance and either persist and support the tumor microenvironment, or escape and proliferate. What generates these diverse populations is unclear. We investigated this in preneoplastic zebrafish livers where the epigenetic regulator, UHRF1, is overexpressed in hepatocytes. Double strand breaks, DNA methylation repatterning, retrotransposon expression, cell cycle withdrawal and activation of Atm and Tp53 dependent senescence were early responses to UHRF1 overexpression. This evolved to generate diverse populations of senescent cells, some which expressed immune and senescence signatures plus anti-apoptotic markers, and others co-expressed proliferative genes. The fate of these populations was dictated by UHRF1 levels and Tp53, as Tp53 loss enabled proliferation of cells with reduced UHRF1 expression but not in cells expressing high UHRF1. The senolytic Navitoclax targeted only a subset of senescent cells. Thus, the diversity of senescent cells driven by epigenetic changes can generate divergent outcomes.

**Short summary:** UHRF1 overexpression in zebrafish hepatocytes induces DNA damage leading to Atm–tp53–dependent senescence with UHRF1 levels dictating whether hepatocytes remain senescent, escape, or undergo senolytic elimination.

## Introduction

Tumor suppression is achieved through the coordination of cell death, senescence and immune clearance of pre-cancerous cells. Cancer arises when damaged cells evade these safeguards, acquiring the ability to exploit resources, withstand stress imposed by the microenvironment, and proliferate. Senescence was originally conceived as a homogenous cellular state characterized by irreversible cell cycle arrest resulting from replicative exhaustion, aging, genotoxic stress or other extreme forms of irreparable damage. It is now clear that the senescence is much more complex phenotype (Avelar *et al*, 2020; Chan & Narita, 2019; Kwiatkowska *et al*, 2023; Ogrodnik *et al*, 2024). Some senescent cells can simultaneously activate tumor suppressive processes, such as cell cycle arrest, and tumor promoting features that reinforce the tumor ecosystem, acquiring mechanisms to either outcompete their neighbors for resources or kill them off (Vishwakarma & Piddini, 2020), and survive in conditions that are suboptimal for non-transformed cells (Chan *et al*, 2024; Ciriello *et al*, 2024; Hausser & Alon, 2020; Loukas *et al*, 2023). There is an incomplete understanding of senescence pleiotropy and of the features that make some senescent cells tumor suppressive or tumor promoting.

While all senescent cells have some features in common, including cell cycle withdrawal and phenotypic changes, their identity and fate vary depending on the senescence inducing stimuli, cellular context and duration of senescence (Campisi & d’Adda di Fagagna, 2007; Chan & Narita, 2019; Ogrodnik *et al*., 2024). DNA damage is a common senescent trigger, and a persistent DNA damage response (DDR) with activation of Tp53, Cdkn1a (p21) and other downstream targets that block proliferation is a feature of most senescent cells (Di Micco *et al*, 2011; Fumagalli *et al*, 2014; Ogrodnik *et al*., 2024). Other defining features are irreversible cell cycle arrest, increased senescence associated beta-galactosidase (SA-β-gal) staining, activation of a pro-inflammatory senescence-associated secretory phenotype (SASP) and senescence associated chromatin alterations, including senescence-associated heterochromatin foci (SAHF) and DNA methylation changes. Not all senescent cells display all these features, reflecting the diversity of senescent cell phenotypes (O’Sullivan *et al*, 2024; Ogrodnik *et al*., 2024).

This diversity translates to distinct fates. Some senescent cells are immunogenic and cleared by the immune system. Others have an opposing phenotype, and become resistant to immune clearance and to apoptosis, persisting even as neighboring cells are cleared (Chan *et al*., 2024; Colucci *et al*, 2025; Kang *et al*, 2011). Persistent senescent cells are dangerous, as some acquire a phenotype that retains the capacity to proliferate when Tp53 or other cell cycle inhibitors fluctuate or are inactivated, allowing these damaged cells to escape, transform and contribute to cancer (O’Sullivan *et al*., 2024; Reyes *et al*, 2018). Other senescent cells, such as those in deep senescence, are incapable of proliferating but are still dangerous, as they support the tumor microenvironment by stimulating angiogenesis or by directly supporting neighboring cancer cells with mitogens, pro-survival signals or factors promoting epithelial to mesenchymal transition (Colucci *et al*., 2025).

Senescent cell diversity is directly relevant to clinical applications, as approaches to selectively reinforce tumor-suppressive senescence while eliminating pro-tumorigenic senescent cells requires methods to identify these different identities. An example is provided by senolytics, such as Navitoclax, which dismantle proteins and pathways that render senescent cells resistant to apoptosis (Colucci *et al*., 2025; Zhu *et al*, 2016). However, Navitoclax is not effective on all senescent cell phenotypes. A Navitoclax pro-drug showed promise in treating a mouse model of hepatocellular carcinoma (HCC) (Yang *et al*, 2025), but it failed in clinical trials for advanced HCC (Emiloju *et al*, 2024). This highlights how senescence targeting therapies must be directed at the population cells that are tumor promoting.

To understand diverse senescent cell phenotypes, it is important to understand how they are generated. Epigenetic changes occur both as a consequence and as a cause of senescence (Cruickshanks *et al*, 2013; Di Micco *et al*., 2011; Hernandez-Segura *et al*, 2018; Ogrodnik *et al*., 2024; Parry *et al*, 2018). While the finding that different senescence stimuli generate distinct patterns of DNA methylation changes (Kwiatkowska *et al*., 2023; Lowe *et al*, 2016) suggests that methylome changes occur as a result of the senescence process, other studies show that manipulation of the DNA methylation machinery causes DNA methylation loss and induces senescence (Chen *et al*, 2025; Jung *et al*, 2017; Yang *et al*, 2017). The widespread loss of DNA methylation on transposable elements (TEs) and the hypermethylation of some promoters, are hallmark features of cancer (Hansen *et al*, 2011; Lister *et al*, 2009) and are already present in senescent cells (Cruickshanks *et al*., 2013; Hlady *et al*, 2017). This shows that modifying the methylome can trigger senescence, and that oncogenic stimuli can cause cancer-promoting changes in the methylome even during the tumor suppressive phase of tumorigenesis.

The epigenetic regulator ubiquitin, PHD and ring finger domain containing protein 1 (UHRF1) regulates the maintenance of DNA methylation during DNA replication (Bostick *et al*, 2007; Qian *et al*, 2008; Sharif *et al*, 2007; Yamaguchi *et al*, 2024), recruits histone methyltransferases that deposit repressive histone modifications (Kong *et al*, 2019; Mancini *et al*, 2021), and is required for DNA repair (Mancini *et al*., 2021; Zhang *et al*, 2016). Our work showing that overexpression of human UHRF1 in zebrafish hepatocytes *Tg(fabp10a:hUHRF1-EGFP)* causes liver cancer (Mudbhary *et al*, 2014) provided functional evidence that an epigenetic regulator can be an oncogene. UHRF1 is upregulated in nearly every type of cancer (Kim & Benavente, 2024), including HCC (Liang *et al*, 2015; Liu *et al*, 2017; Mudbhary *et al*., 2014; Wang *et al*, 2023). We previously showed that within 2 days of hUHRF1 overexpression in zebrafish hepatocytes, classical markers of senescence, including *tp53*, SA-β-gal and cell cycle withdrawal were present, with liver cancer developing 2 weeks later (Ajouaou *et al*, 2023; Mudbhary *et al*., 2014). Here, we used this system to investigate the mechanism of UHRF1-induced senescence and found that this single oncogene can induce pleotropic senescent cell identities that evolve over time. Within hours of hUHRF1 overexpression, hepatocytes withdraw from the cell cycle, activate TEs and have DNA double strand breaks (DSBs), activating ATM and Tp53 to promote some, but not all, features of senescence. As senescence progresses, distinct populations of senescent cells appear which are distinguished by hUHRF1 levels, tp53 activation and mitogen responsiveness. At later stages, cells with reduced hUHRF1 levels can proliferate when Tp53 is removed, whereas cells with the highest hUHRF1 expression cannot. Only some cells are sensitive to Navitoclax, allowing expansion of other populations. This suggests that in UHRF1-driven cancers, senolytic treatment may inadvertently promote tumor growth and highlight the importance of understanding senescence heterogeneity in cancer progression.

## Results

### UHRF1 overexpression rapidly induces a DDR and SASP

The *fabp10a* promoter that drives hUHRF1-EGFP in hepatocytes is highly expressed in hepatocytes as soon as they differentiate (Her *et al*, 2003). A control transgene under this promoter, *Tg*(*fabp10a:nls-mCherry*), can be detected in all larvae starting at 72 hours post-fertilization (hpf), whereas EGFP can be detected in 100% of *Tg(fabp10a:hsa.UHRF1-EGFP)^mss1a^*larvae (hereafter called *hUHRF1*) only at 80 hpf and later (Figure 1A). We reported cell cycle withdrawal at 120 hpf and strong SA-β-gal staining in nearly all *hUHRF1* livers at 120 hpf, but not at 96 hpf (Mudbhary *et al*., 2014). We asked when hUHRF1 overexpression induced cell cycle withdrawal in the liver using EdU incorporation. At 80, 96 and 120 hpf, there were 25%, 15% and 2% EdU positive cells in controls, respectively, which were significantly reduced to 15% and 5% in *hUHRF1* larvae at 80 and 96 hpf, respectively. There was no difference in EdU incorporation at 120 hpf, since nearly all cells in control livers had completed S phase at this stage (Figure 1B). Using nuclear morphology to identify hepatocytes showed that both hepatocytes and non-hepatocytes incorporate EdU in control livers at 96 hpf, but virtually no hepatocytes incorporated EdU in *hUHRF1* livers (Figure 1C).

**Figure 1.**
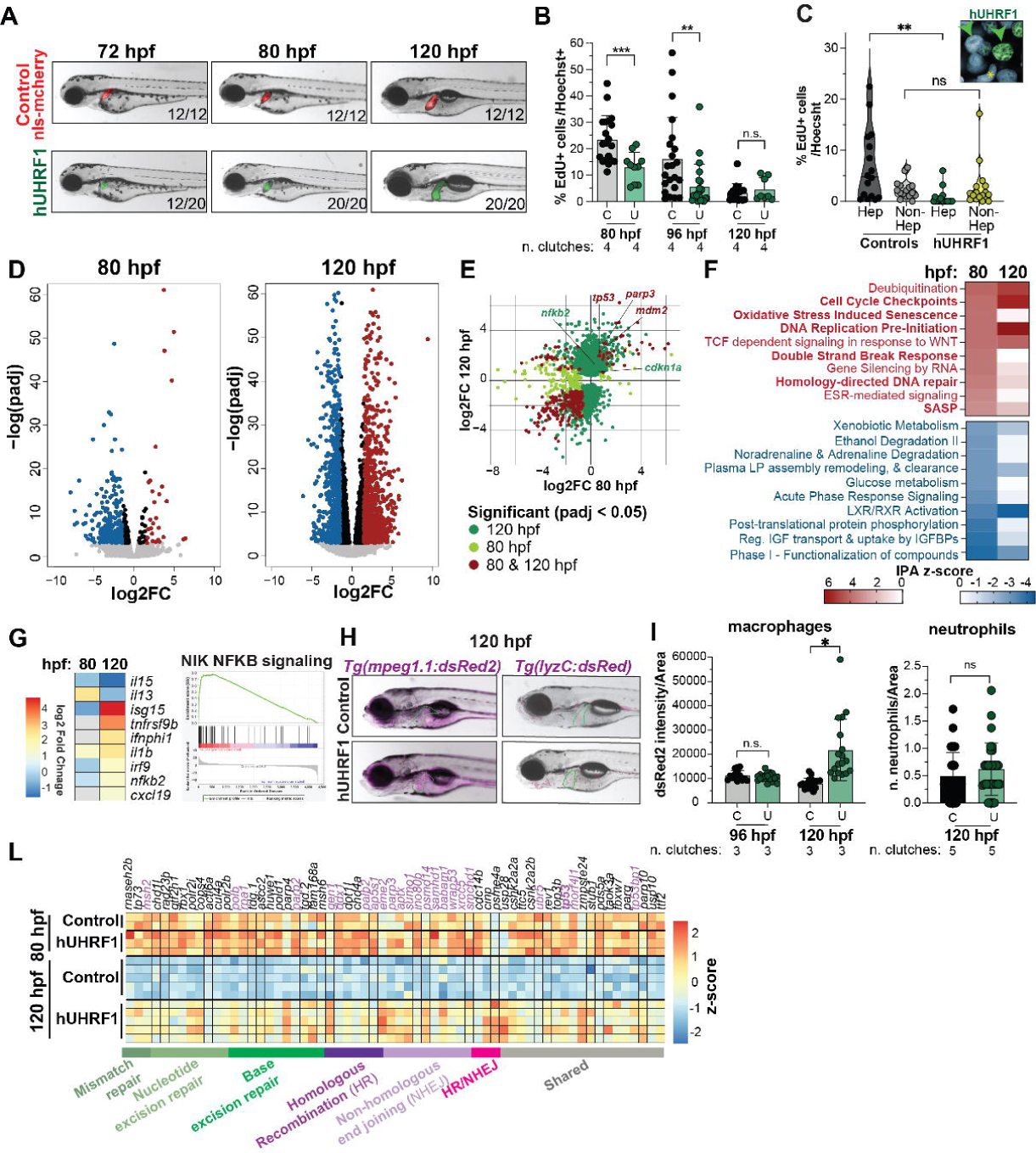
UHRF1 overexpression rapidly induces a DDR and SASP. **A.** Images of live *hUHRF1* and control *Tg(fabp10a:nls-mCherry)* larvae at 80, and 120 hpf. Numbers indicate the number of larvae with fluorescent livers at each time point. **B.** EdU incorporation in *hUHRF1* and control livers at 80, 96, and 120 hpf normalized to the total number of nuclei in each sample. *** p-value < 0.001, ** p-value < 0.01 by unpaired t-test. **C.** EdU positive cells in 96 hpf livers were scored as hepatocytes based on round nuclear morphology (green arrow in inset) and non-hepatocytes with non-spherical nuclei (* in inset). ** p-value < 0.01 by unpaired t-test. **D.** Volcano plot of differential expressed genes of *hUHRF1* compared Controls livers at 80 and 120 hpf. **E.** Crossplot of the Log2FC of differentially expressed genes in *hUHRF1* livers at 80 and 120 hpf. **F.** IPA of the top 10 most upregulated (up) and downregulated pathways (down) in *hUHRF1* livers at 80 hpf and their relative changes in *hUHRF1* livers at 120 hpf. **G.** Heatmap of Log2 fold change of a subset of genes associated with SASP at 80 and 120 hpf and the Gene Set Enrichment plot of the NFKB pathway identified from RNAseq analysis of 120 hpf livers from *hUHRF1* compared to controls. **H.** Representative images of macrophages (purple) in *Tg(mpeg1.1:dsRed2)* and neutrophils (purple) in *Tg(lyz:dsRed)* in *hUHRF1* and control larvae at 120 hpf. Liver is outlined in green. **I.** Quantification of macrophage area and number of neutrophils recruited in the liver at 96 and 120 hpf. * p-value < 0.05 by unpaired t-test. **L.** Heatmap of genes involved in DNA damage repair at 80 and 120 hpf.

We eliminated the possibility that the transgene insertion caused senescence due to a bystander effect by showing that CRISPR-Cas9 mutagenesis of the transgene (Figure S1A-B) rescued SA-β-gal staining (Figure S1C). We mapped the insertion of the transgene to Chromosome 14 and used analyzed bulk RNAseq data from livers from *hUHRF1* and control larvae at 80 hpf and 120 hpf (Table S1) to examine the expression of genes at this locus (Figure S1D), showing none were altered in *hUHRF1* livers at 80 or 120 hpf (Figure S1E), eliminating the likelihood of a positional effect of the transgene.

To determine the immediate and sustained transcriptional response to hUHRF1 overexpression, we assessed gene expression at 80 hpf, the earliest time point when hUHRF1-EGFP is detected in 100% of larvae (Figure 1A) and the first functional response is observed (Figure 1B) compared to the time when the senescence phenotype is fully penetrant, characterized by robust SA-β-gal and activation of *tp53* and its downstream targets (Ajouaou *et al*., 2023; Mudbhary *et al*., 2014). At 80 hpf, there were 445 significantly differentially expressed genes (DEGs; p-value adjusted < 0.05), with 51 upregulated and 394 downregulated compared to 5,353 genes at 120 hpf (p-value adjusted < 0.05), with 2640 upregulated and 2713 downregulated (Figure 1D, Table S1). Importantly, most of the genes from 80 hpf were similarly expressed at 120 hpf, including *tp53* and its key target genes (Figure 1E). Ingenuity Pathway Analysis (IPA) identified pathways involved in SASP, cell cycle checkpoints and the response to double stranded breaks (DSB) induced as early as 80 hpf and sustained through 120 hpf (Figure 1F), while downregulated pathways at both time points reflected a range of hepatocyte functions (Figure 1F). Interestingly, however, there was differential enrichment of some pathways at these time points, suggesting an evolution of the senescence phenotype. For instance, many SASP genes and a key upstream immune regulator, NF-kB, were induced only modestly or not at all at 80 hpf but were highly induced at 120 hpf (Figure 1G), reflecting progression of the senescence phenotype. Transgenic reporter lines of macrophages, *Tg(mpeg1.1:dsRed),* and neutrophils, *Tg(lyzC:dsRed),* showed that the number of macrophages in the liver were significantly increased at 120 hpf but not at 96 hpf while neutrophils were not increased either at 96 or 120 hpf (Figure 1H-I), suggesting that SASP and immune recruitment is a delayed response to hUHRF1 overexpression. We next examined the DDR in *hUHRF1* livers and found that genes involved in multiple DDR pathways, in particular homologous repair and non-homologous end joining were induced by hUHRF1 overexpression at both 80 and 120 hpf (Figure 1L). This included *tp53*, *atm* and downstream targets which we previously showed were highly upregulated at 120 hpf in *hUHRF1* livers (Ajouaou *et al*., 2023; Mudbhary *et al*., 2014). These data suggest that DNA damage and cell cycle withdrawal is an early response to hUHRF1 overexpression, that it evolves over time to induce SASP and macrophage recruitment.

### DSBs and TE activation associated with DNA methylation loss are early responses to hUHRF1 overexpression

These data suggested that hUHRF1 overexpression induced DNA damage. To assess this, we stained for γH2AX foci, as marker of double strand breaks (DSBs), and Phosphorylated replication protein A (p-Rpa), as it accumulates on single stranded DNA during replication stress or DSBs repair. γH2AX foci were absent from control nuclei at 80 or 96 hpf but were detected in *hUHRF1* samples as early as 80 hpf and persisted to 96 hpf (Figure 2A). p-Rpa was detected in *hUHRF1* samples at 96 hpf but not at 80 hpf (Figure 2B). This indicates that UHRF1 overexpression causes DSBs followed by formation of single stranded DNA, either as a byproduct of DNA repair or possibly as a response to replication stress.

**Figure 2.**
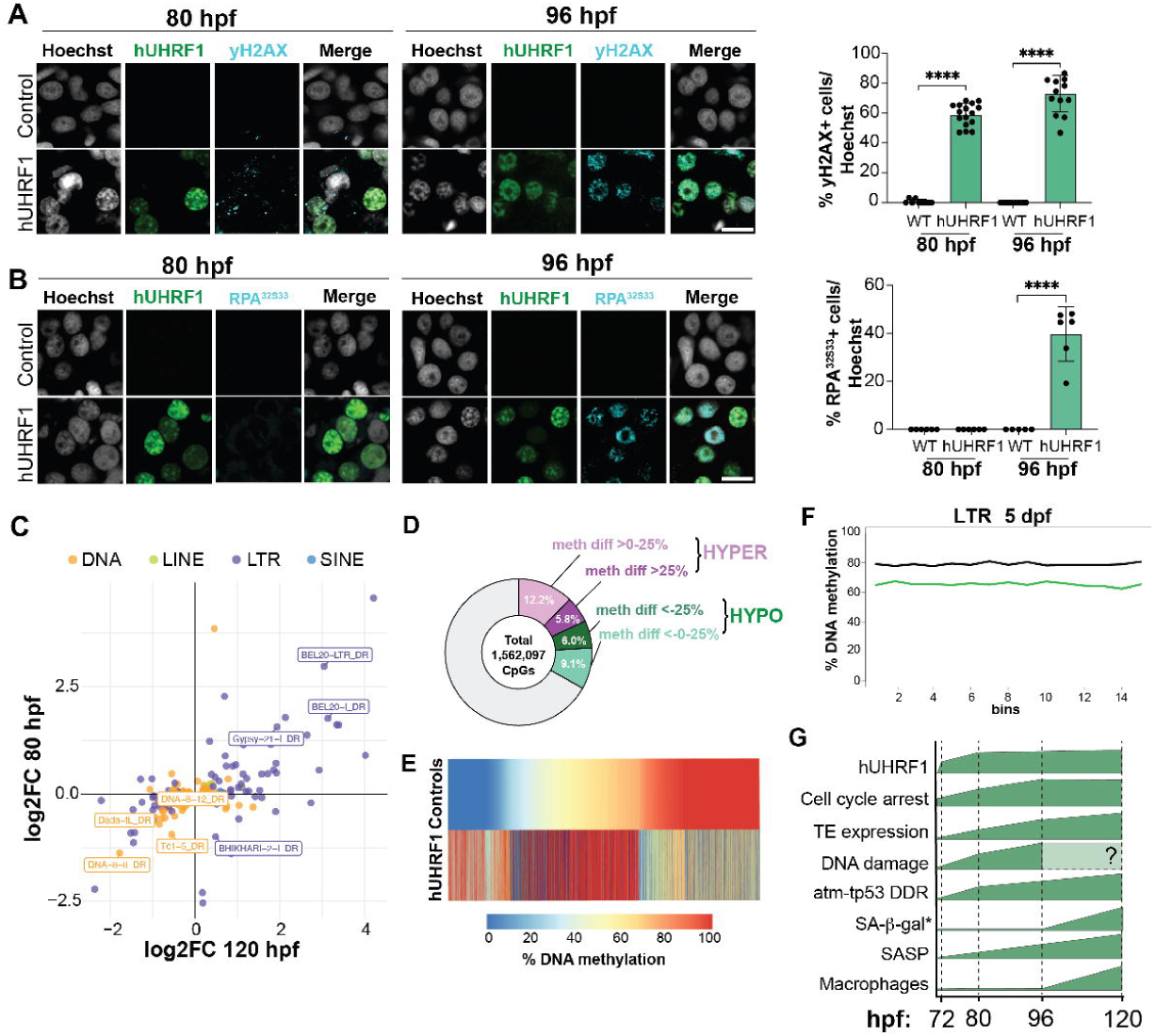
DSBs and TE activation associated with DNA methylation loss are early responses to hUHRF1 overexpression. Immunofluorescence of γH2AX (**A**) and p-Rpa (**B**) staining of *hUHRF1* and control livers at 80 and 96 hpf. Quantification of the percentage of positive nuclei in the liver for each marker. **** p-value < 0.0001 by unpaired t-test. **C.** Crossplot of Log2 fold change of differentially expressed TEs in *hUHRF1* and control livers at 80 and 120 hpf. **D.** Pie chart classification of 1,562,097 CpGs 10X covered by RRBS in all samples at 120 hpf. CpGs were categorized as hypermethylated or hypomethylated by an FDR <0.01 and then classified as 0-25% change or >25% change in *hUHRF1* livers compared to controls. **E.** Heatmap of CpGs that were significantly differentially methylated (FDR <0.01) and had an effect size of >25% change rank ordered by level of methylation in controls. **F.** Metaplot of CpG methylation levels on all LTR transposons in *hUHRF1* (green) and control (black) livers at 120 hpf. **G.** Time course of senescence associated responses caused by hUHRF1 overexpression in hepatocytes 80, 96 and 120 hpf. * SA-b-gal data are extracted from previous publication (Mudbhary *et al*., 2014).

TEs can cause DSBs (Gasior *et al*, 2006) and since UHRF1 is required for DNA methylation (Bostick *et al*., 2007; Sharif *et al*., 2007), which is a primary mechanism of TE suppression, we investigated TE activation in bulk RNAseq of *hUHRF1* livers. LTR retrotransposons were induced at both 80 hpf and 120 hpf (Figure 2C), indicating that the epigenetic mechanisms suppressing these TEs are deregulated as early as 80 hpf. We next profiled DNA methylation at 120 hpf which was the earliest time point when we could obtain sufficient genomic DNA from pools of dissected livers to carry out low input reduced representation bisulfite sequencing (RRBS). We obtained 10X coverage of over 1.5 million CpGs in all samples analyzed. hUHRF1 overexpression caused a change in methylation levels in nearly 1/3 of all CpGs analyzed; with 15% of all CpGs having lost methylation (Figure 2D; Table S2). Loss of methylation was most pronounced on the CpGs that were fully methylated in controls, and the majority of CpGs that gained methylation were detected as partially methylated in controls (Figure 2E). Importantly, LTR retrotransposons were hypomethylated in *hUHRF1* livers (Figure 2F), reminiscent of the profile in zebrafish mutants where DNA methylation is depleted (Chernyavskaya *et al*, 2017; Magnani *et al*, 2021). This suggests that DNA hypomethylation of retrotransposons in *hUHRF1* livers induces their expression and we speculate this as a source of DNA damage in this model. Together, these data define the temporal sequence of responses to hUHRF1 overexpression, with TE activation, DNA damage, DDR activation and cell-cycle arrest occurring as immediate response, followed by SASP, macrophage recruitment and SA-β-gal positive staining within 2 days (Figure 2G). We hypothesize that these events are a downstream effect of DNA methylation changes caused by UHRF1 deregulation during S-phase.

### The *atm*-*tp53* pathway is required for distinct UHRF1-induced senescence features

We previously identified *tp53*, *atm* (Ajouaou *et al*., 2023) and Tp53 downstream targets to be induced at 120 hpf, and that heterozygosity for *tp53* reduced SA-β-gal staining and accelerated tumor onset in *hUHRF1* larvae (Mudbhary *et al*., 2014). We thus investigated the role of Tp53 activators, ATM and ATR in hUHRF1-mediated senescence. Homozygous loss of function mutation in *tp53* (Berghmans *et al*, 2005) or *atm* (Ajouaou *et al*., 2023) abrogated SA-β-gal staining in *hUHRF1* larvae (Figure 3A), while inhibiting ATR using a pharmacological inhibitor (VE-821; Figure S2) did not reduce SA-β-gal staining alone or in combination with *atm* mutation (Figure 3A). This indicates that both Atm and Tp53, but not Atr, are required for hUHRF1-induced senescence, and that Atr does not compensate for *atm* loss in this setting.

**Figure 3.**
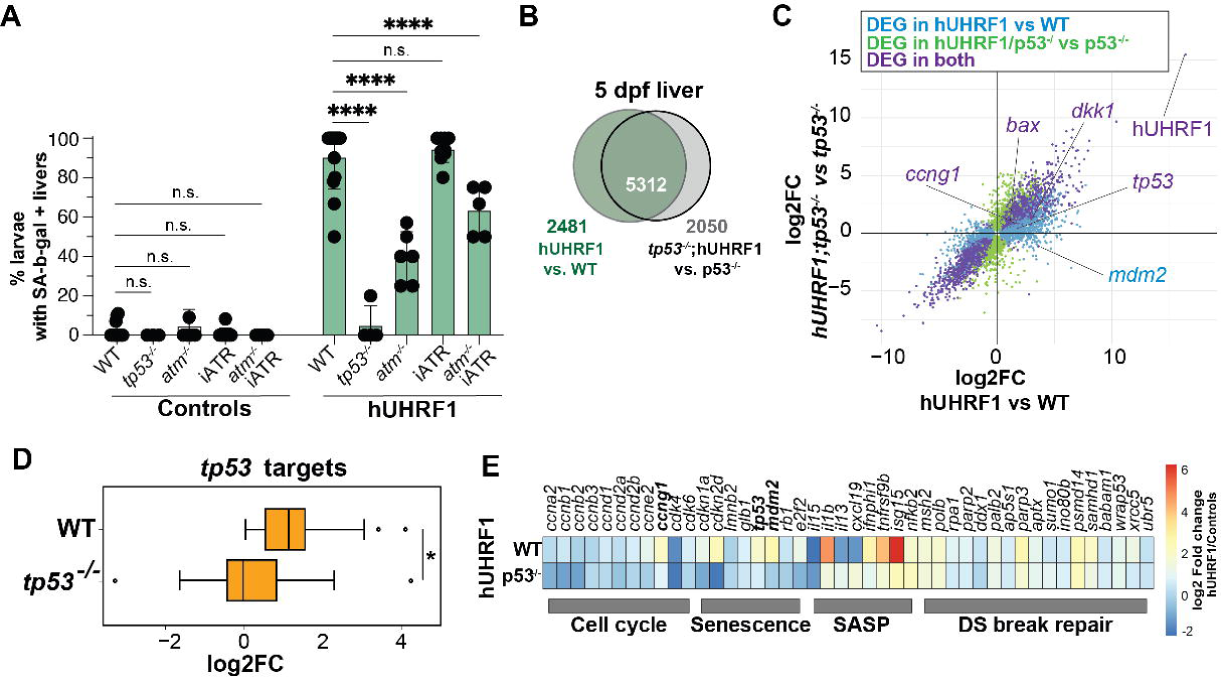
The *atm*-*tp53* pathway is required for distinct UHRF1-induced senescence features. **A.** Percent of SA-β-gal positive livers at 120 hpf in *hUHRF1* and control livers in a WT background or mutation of *tp53* or *atm*, with chemical inhibition of Atr using 10 μM VE821 (iATR), or combination of *atm* mutation and 10 μM VE821. **** p-value < 0.0001 by unpaired t-test. Each dot represents the percent of positive larvae in a clutch, with at least 10 larvae scored per clutch. **B.** Venn diagram of DEGs in the liver of *hUHRF1* compared WT or in *hUHRF1;tp53^-/-^* compared *tp53^-/-^.* **C.** Crossplot of Log2FC of differentially expressed genes in *hUHRF1* compared to WT controls or *hUHRF1*;*tp53^-/-^* to *tp53^-/-^* at 120 hpf. **D.** Box plot of the Log2 fold change of *tp53* target genes (Fischer, 2017) upregulated in *hUHRF1* on a WT background (WT) compared to the expression of the same genes in *hUHRF1*;*tp53^-/-^* compared to *tp53^-/-^* (*tp53^-/-^)*. **E.** Heatmap of of select Tp53 target genes relevant to senescence in *hUHRF1* on a WT background (WT) compared to the expression of the same genes in *hUHRF1*;*tp53^-/-^* compared to *tp53^-/-^* (*tp53^-/-^)*.

To investigate the *tp53*-dependent senescence phenotypes induced by hUHRF1 overexpression, we carried out bulk RNA-seq on control and *hUHRF1* livers with either wild type or *tp53^-/-^* mutant background (Figure 3B; Table S3). There were 2,760 DEGs in *tp53^-/-^* livers compared to WT (p-value adjusted < 0.05; Figure S3A, Table S3), with deregulated pathways representing a range of cellular functions including upregulation of diverse signaling pathways and down regulation of metabolic processes (Figure S3B). The majority of the differentially expressed genes in *hUHRF1* were similarly expressed in *hUHRF1;p53^-/-^* samples (Figure 3B-C), with the exception of canonical *tp53* target genes (Fischer, 2017) such as *mdmd2*, *bax*, *dkk1*, *ccng1* and *gadd45ab* (Figure 3C-D, S3C). Importantly, genes involved in senescence and SASP were mostly rescued in *tp53^-/-^* mutants while genes involved in apoptosis (Figure S3C) and many genes in the DDR pathway remained largely unchanged (Figure 3E). Notably, all the genes that drive cell division remain suppressed in *hUHRF1* livers, even when *tp53* was removed (Figure 3E; Table S3). Together, these data indicate that hUHRF1 overexpression causes DSBs and activates the ATM-Tp53 pathway, which in turn induces senescence features including SA-β-gal activity and the SASP. However, at this early stage of senescence, *tp53* loss is insufficient to allow UHRF1-overexpressing cells to re-enter the cell cycle, as seen in other models (Reyes *et al*., 2018).

### UHRF1 overexpression in hepatocytes changes liver composition and hepatocyte function

To identify cell type-specific gene expression changes caused by hUHRF1 overexpression, we performed single cell RNAseq (scRNAseq) on pooled livers. These were manually dissected from 120 hpf *hUHRF1* and control larvae expressing membrane-targeted EGFP under the same hepatocyte promoter (*Tg(fabp10a:CAAX-EGFP)*; hereafter referred to as Controls; Figure 4A). This time point was chosen because it was the stage at which all senescence markers were present and a sufficient number of liver cells could be isolated for sequencing. After quality control, the combined dataset contained 23,121 cells segregated into 10 clusters, each assigned to different hepatic cell type according to expression of established cell identity markers (Morrison *et al*, 2022; Oderberg & Goessling, 2023) (Figure 4B-C; Table S4). There were 4 populations of hepatocytes and, as expected, together these 15,986 cells constituted the majority of all cells (69.1% of total). Biliary epithelial cells (BEC), characterized by the expression of *alcama*, *sox9b*, *epcam* and keratin genes, represented the next largest population of cells in the combined dataset (2,216 cells; 9.2% of total), followed by endothelial and mesenchymal cells, which were divided into 2 populations, hepatic stellate cells (HSC) and fibroblasts based on canonical markers (Oderberg & Goessling, 2023) and showed distinct and overlapping gene sets reflecting their function (Figure 4B-D). The hUHRF1 and EGFP sequences were added to the danRer10 genome to allow transgene mapping. In both samples, the transgene was expressed primarily in hepatocyte populations (Figure S4B-C).

**Figure 4.**
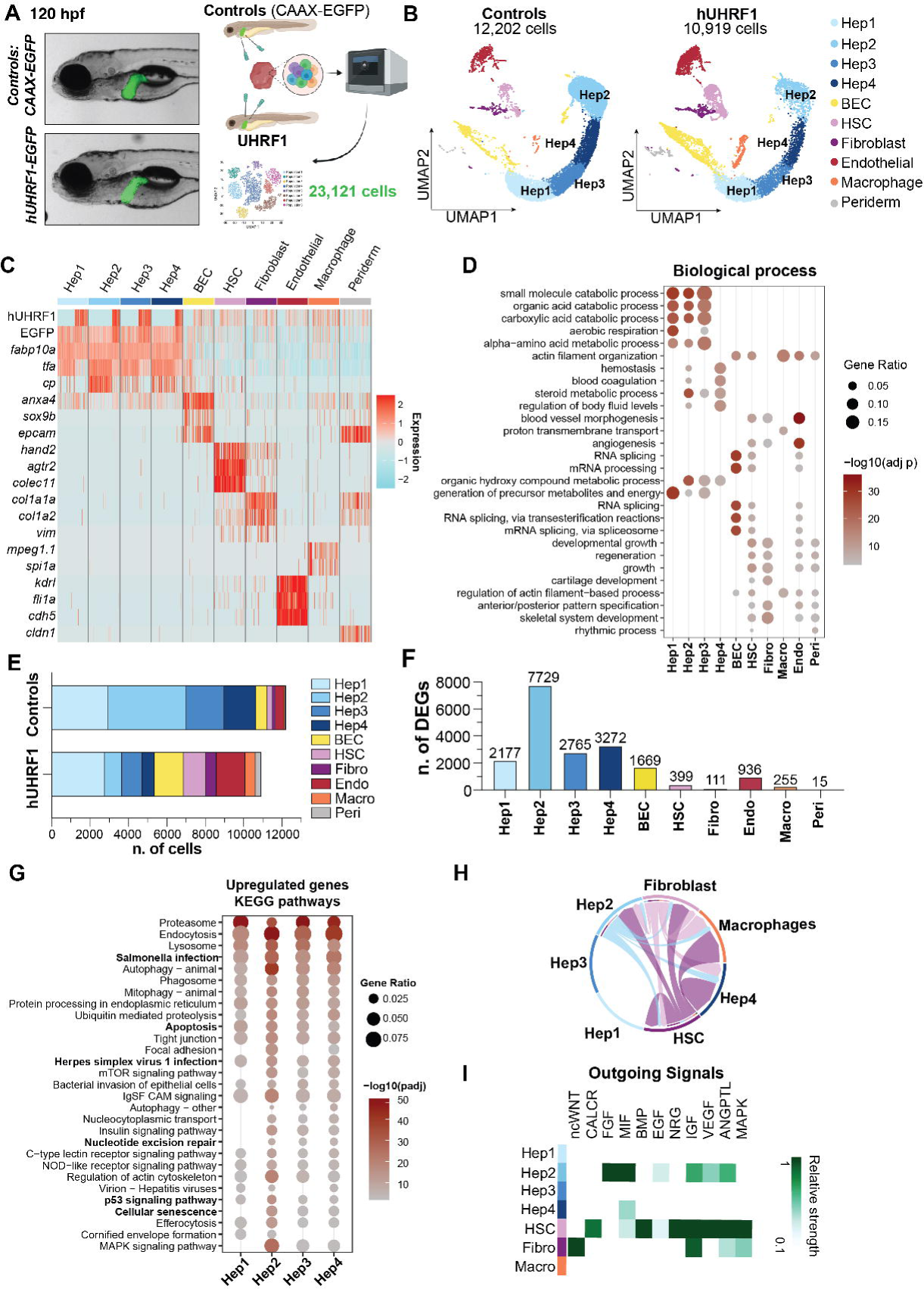
UHRF1 overexpression in hepatocytes changes liver composition and hepatocytes function. **A.** Scheme of the approach and representative images at 120 hpf of samples used for scRNAseq analysis. **B.** UMAP plot of cells analyzed by scRNAseq of *hUHRF1* and control livers. **C.** Heatmap of identity markers in hepatic cell populations pooled for controls and *hUHRF1* samples. **D.** Gene Ontology of enriched genes (p-value adjusted < 0.05 & Log2FC > 0.5) in each cell population from the combined dataset. **E.** Number of cells in each population in *hUHRF1* and control livers. **F.** Number of DEGs (padj < 0.05) in *hUHRF1* compared to controls in each population. **G.** KEGG pathways analysis of DEGs (padj < 0.05 & Log2FC > 1.5) in *hUHRF1* compared to controls for each population of hepatocytes. **H.** Chord diagram showing of cell-cell communication, with the width of each chord representing the proportion of cells with that signal and the arrows representing the direction of the signal. **I.** The top outgoing signaling molecules are inferred based on the strength of the signaling in each population.

Gene Ontology analysis of significantly enriched genes in each population (p-value adjusted < 0.05 & Log2 Fold Change > 0.5) identified hepatocyte populations with different functions, with Hep4 distinguished from others by pathways associated with hemostasis and blood coagulation and Hep1-3 were enriched in metabolic processes (Figure 4D). This indicates that, as in mammals (Aizarani *et al*, 2019) different populations of hepatocytes in zebrafish assume distinct functions.

There was a striking shift in the distribution of cell populations between controls and *hUHRF1* samples (Figure 4E), with depletion of hepatocyte populations and relative expansion of non-parenchymal cells in *hUHRF1* samples (Figure 4B, E). There was a marked change in the relative distribution of hepatocyte populations, except for Hep1, which was equally abundant in controls and *hUHRF1* samples (Figure 4B, E). Hep2 was the most depleted population in *hUHRF1* samples, and this population had the greatest number of DEGs (Figure 4E-F). Analysis of KEGG pathways from the DEGs identified in each hepatocyte population compared to the same population in controls (p-value adjusted < 0.05 & Log2FC > 1.5) showed that all hepatocyte populations in *hUHRF1* samples were enriched for Tp53 signaling, apoptosis and those related to infection (Figure 4G). In contrast, Hep2, Hep3 and Hep4 populations were enriched for pathways associated with cellular senescence and both pro-apoptotic and proliferative components of the MAPK signaling pathway (Figure 4G-H). The expansion of macrophages in the *hUHRF1* dataset (Figure 4B, E) supports imaging data showing macrophage infiltration in the liver at this time point (Figure 1I). It is not clear whether the increase in representation of mesenchymal cells could represent collection bias or expansion of these populations in *hUHRF1* samples.

Next, we performed cell-cell communication analysis among hepatocyte populations in *hUHRF1* samples to determine potential functional differences in these cell types using CellChat. Interestingly, the most reciprocal signaling occurred between Hep2 and mesenchymal cells, while Hep4 and macrophages were only involved in receiving signals (Figure 4H). We investigated the specific signaling pathways involved in each cell type and found that Hep2 cells were enriched for Angptl and Mif, two important signaling pathways in SASP, VEGF, which is important for endothelial outgrowth and the tumor microenvironment, and for IGF, FGF, and EGF, all mitogenic signals (Figure 4I). This demonstrates that 2 days of hUHRF1 overexpression generates major changes in hepatic cell composition and diversifies the hepatocyte cell populations.

### UHRF1 overexpression causes pleiotropic senescence phenotypes

To better define the senescence phenotypes caused by hUHRF1 overexpression we assessed genes involved in senescence across populations and found that they were enriched in Hep2 (Figure 5A). The senescence associated genes identified as affected by *hUHRF1* overexpression in bulk RNAseq (Figure 3E) were all enriched in Hep2 (Figure 5B), but both *tp53* and key downstream targets, *cdkn1a* and *mdm2* were highly expressed in all hepatocytes, as was the key SASP regulator, *nfkb2* and multiple DSB repair genes (Figure 5B). This indicates that some markers of senescence phenotype were present in all hepatocytes at this time point, but that there was a clear distinction of one population of hepatocytes that had additional features. For instance, Hep2 cells showed high expression of cell cycle genes (Figure 5B) consistent with this population of senescent cells having high signaling sending and receiving capacity.

**Figure 5.**
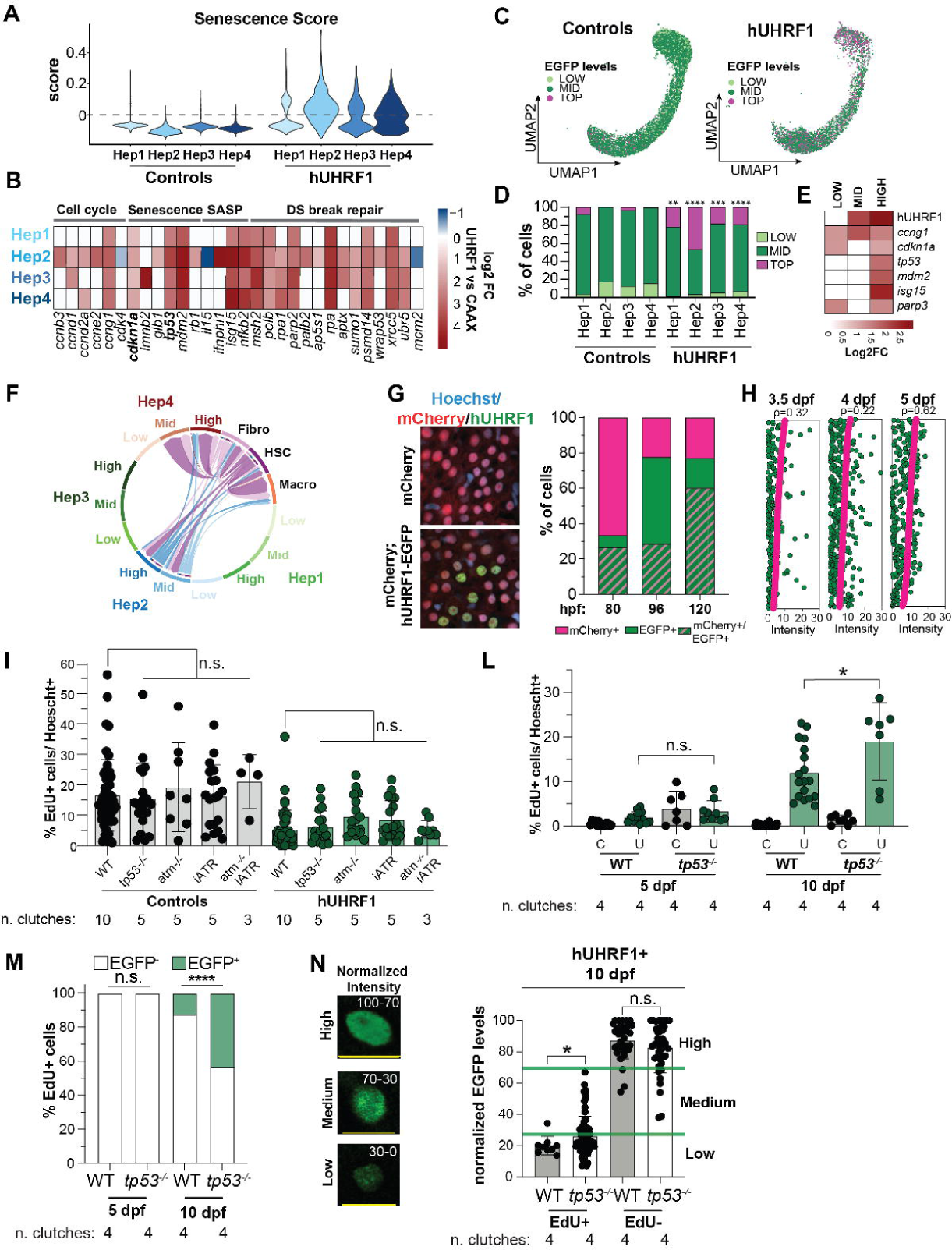
UHRF1 overexpression causes pleiotropic senescence phenotypes. **A.** Violin plot of the enrichment score calculated on all genes in the KEGG Senescence pathway in hepatocytes. **B.** Heatmap of key genes involved in senescence with Log2FC calculated from *hUHRF1* compared Controls for each population. **C.** UMAP of hepatocytes divided in high (top 10 % EGFP expressing cells), mid and low (bottom 10 % EGFP expressing cells) based on EGFP expression in *hUHRF1* and control livers. **D.** Percent of cells in each population that have high, mid or low expression of EGFP. ** p-value < 0.01, *** p-value < 0.001, **** p-value < 0.0001 by Chi-square test on percentage of high cells in *hUHRF1* compared controls for each population. **E.** Heatmap of a panel of genes involved in Senescence. Log2FC is calculated as enriched genes in each category for *hUHRF1* compared to Controls. **F.** Chord diagram of cell-cell communication of Hep populations divided in high, mid low EGFP expression with mesenchymal cells and macrophages. The width of each chord representing the proportion of cells with that signal and the arrows representing the direction of the signal. **G.** Representative images of hUHRF1-EGFP and nls-mCherry in controls and *hUHRF1* at 120 hpf. Quantification of cells positive for EGFP and mCherry, EGFP only and mCherry only in *Tg(fabp10a:hUHRF1;nls-mCherry)* and controls (mCherry) livers. **H.** Quantification of scaled fluorescent intensity normalized on maximum intensity for each channel for each liver of EGFP and mCherry in *hUHRF1;mCherry* livers and controls across time points. **I.** EdU incorporation at 96 hpf in *hUHRF1* and control livers in WT background or with *tp53*^-/-^, *atm*^-/-^, 10 µM of VE821 (iATR) alone or combination with *atm*^-/-^. n.s. p-value > 0.05 with unpaired t-test. **L.** EdU incorporation in *hUHRF1*, WT control, *tp53^-/-^;hUHRF1* and *tp53^-/-^* livers at 5, and 10 dpf. * p-value < 0.05 by unpaired t-test. **M.** Percent of EdU positive cells that are either EGFP positive or negative. **** p-value < 0.0001 by Chi-square test. **N.** Relative intensity of EGFP in each cell, normalized for each liver on the most EGFP expressing cells, in *hUHRF1* (WT) and *tp53^-/-^;hUHRF1(tp53^-/-^).* For each panel, number of biological replicates (clutches) is indicated.

A recent study showed that Ras level determines behavior and fate of senescent cells. Cells with high Ras levels are immunogenic, leading to their elimination by the immune system, whereas cells with lower Ras levels are immunoresistant and can escape senescence (Chan *et al*., 2024). We hypothesized that different levels of hUHRF1 could mediate the different senescent states identified in hepatocytes. To test this, we categorized hepatocytes from control and *hUHRF1* samples into clusters with cells with the lowest 10% of EGFP expression classified as low, those in the highest 10% as the high expressing, and the remaining cells assigned as middle (mid; Figure 5C-D). Nearly all the cells in controls livers expressed mid or low levels of EGFP, while cells in *hUHRF1* samples were heterogenous, with Hep2 having the highest proportion of high expressing cells (Figure 5D). High levels of hUHRF1 were associated with deep senescent state, as *cdkn1a*, *tp53* and other senescence associated genes were enriched in the high expressing cells (Figure 5E), and high and mid cells from Hep2 and Hep4 were mostly engaged in cross-talk with each other, mesenchymal cells and macrophages (Figure 5F). In contrast, Hep1, Hep3, and all hepatocyte populations with low hUHRF1 expression showed no detectable cell-cell communication. This indicates that hUHRF1 levels determine distinct senescent-cell phenotypes and regulate their capacity to signal to neighboring hepatocytes.

To examine hUHRF1 heterogeneity and the relationship to function *in vivo*, we used confocal imaging of livers expressing nuclear mCherry in hepatocytes *Tg(fabp10a:nls-mCherry)* together with *hUHRF1-EGFP* and scored each nucleus as positive or negative for either mCherry or EGFP (Figure 5G-H). By 80 hpf most hepatocytes expressed nls-mCherry with minimal cell to cell variability (Figure S5A), and by 120 hpf, the majority of hepatocytes were positive for both transgenes (Figure 5G). However, the levels of EGFP were heterogenous, ranging from negligible to high (Figure 5H). Importantly, EGFP levels were not correlated with nls-mCherry levels (Figure 5H), and *in situ* hybridization detected heterogenous hUHRF1 mRNA expression (Figure S5B) suggesting transcriptional heterogeneity as the cause.

We previously showed that loss of *tp53* causes increased cell proliferation in stages just preceding tumor formation and accelerated tumor formation caused by hUHRF1 overexpression (Mudbhary *et al*., 2014). We hypothesized that the increase in proliferation would only be possible if cells had both acquired the ability to respond to proliferative signals and all cell cycle inhibitors were dismantled. We therefore investigated whether Tp53 signaling acts as a gatekeeper that maintains senescence in cell populations susceptible to proliferative cues, by assessing EdU incorporation in *hUHRF1* livers with *tp53* and *atm* mutation in early senescence (96 hpf). However, despite effectively blocking SA-β-gal staining (Figure 3A), neither restored proliferation at this stage. This indicates that in some stages of senescence, cell cycle arrest is maintained by factors acting upstream of *atm* and *tp53*, potentially as a direct consequence of UHRF1 overexpression.

We next reasoned that the ability for tp53 loss to increase proliferation at later stages would be related to the distinct senescent phenotypes induced by hUHRF1 and correlated with hUHRF1 expression levels. We tested this by assessing which cells could proliferate when *tp53* was removed at 10 dpf. As expected, EdU incorporation increased in *hUHRF1* livers in WT background and nearly all cells incorporating EdU lacked hUHRF1-EGFP (Figure 5 L-M). Removing *tp53* increased EGFP-positive proliferating cells by three-fold (Figure 5M). We hypothesized that the high *tp53* signaling observed in cells with the highest hUHRF1 expression reflected deep senescence, and these would be proliferation incompetent, whereas the cells with lower expression might be able to reenter the cell cycle if *tp53* was removed. To test this, EGFP levels in all cells were measured and categorized as high, medium and low (Figure 5N). We then assessed the level of EGFP in the cells that were EdU positive. In WT samples, all EdU positive cells had either low or medium EGFP levels, whereas in *tp53* mutants, the EdU positive cells had higher EGFP levels. However, cells with the highest EGFP levels were EdU negative regardless of *tp53* presence or loss (Figure 5N). Thus, hUHRF1 levels generates cells with distinct senescent phenotypes and capacities, some which can respond to proliferative signals but are restrained by *tp53*, others which cannot.

### Navitoclax removes one population of senescent cells

We next asked whether the pleiotropic senescent phenotypes caused by hUHRF1 heterogeneity conferred differential sensitivity to senolytic therapy. Analysis of genes targeted by senolytics showed that most of the targets were enriched in Hep2 cells (Figure 6A). This suggested that only a subset of senescent cell populations would be sensitive to senolytic elimination and that Navitoclax would be more effective than Dasatinib. Treatment of hUHRF1 larvae with Navitoclax significantly reduced SA-β-gal staining, whereas Dasatinib had no significant effect (Figure 6B). Navitoclax also increased EdU incorporation in hepatocytes of *hUHRF1* livers (Figure 6C). Together, these data show that senolytic therapy may be effective at removing some, but not all senescent cells, sparing some populations which may be prone to escape senescence and re-enter the cell cycle (Figure 6D).

**Figure 6.**
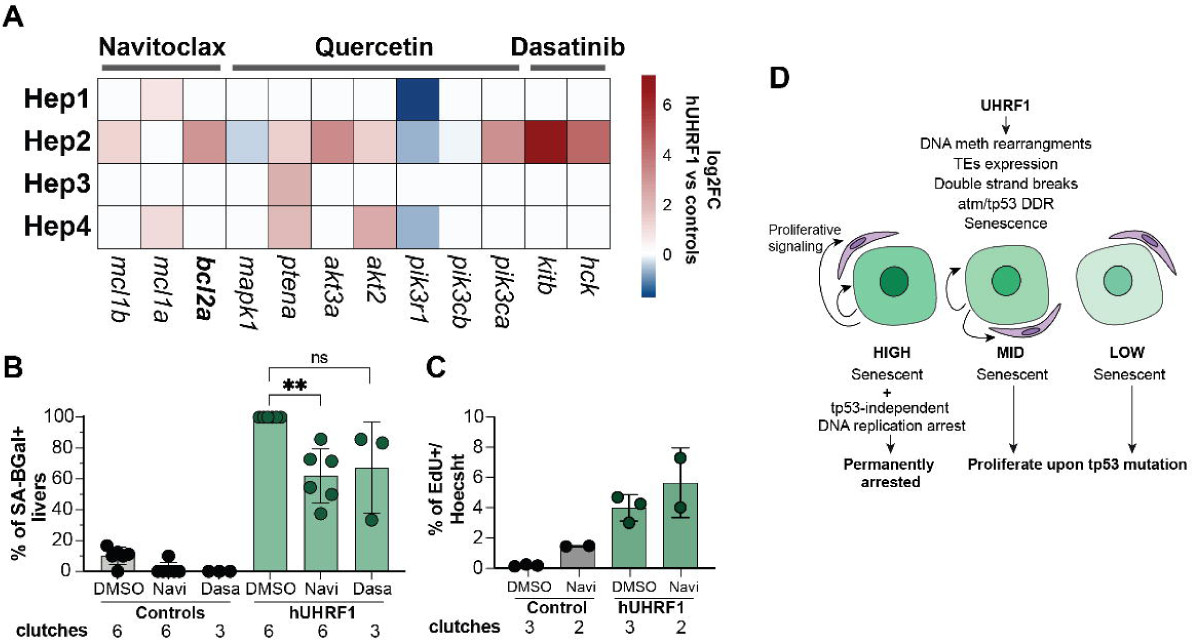
Navitoclax reduces hUHRF1-induced senescence. **A.** Heatmap of Log2FC of senolytic targets in *hUHRF1* hepatocytes based on scRNAseq compared to controls. **B.** SA-β-gal positive livers at 7 dpf in *hUHRF1* and control livers upon treatment with 25 μM Navitoclax, 2.5 μM Dasatinib or DMSO (Vehicle control) from 4 till 7 dpf. ** p-value < 0.01 by unpaired t-test. Each dots represent the percent of positive larvae per clutch. **C.** EdU incorporation at 7 dpf in *hUHRF1* and control livers upon treatment with 25 μM Navitoclax or DMSO from 4 till 7 dpf. **F.** Model of mechanism generating pleiotropic senescent phenotypes induced by UHRF1 overexpression.

## Discussion

The cause, behavior, and fate of cells with different senescent phenotypes is an important, unanswered area in cancer biology. Moreover, it is not clear how a single oncogene can induce diverse cell identities and fates. We address this using a precancer model in zebrafish whereby the epigenetic regulator and oncogene, UHRF1 is overexpressed in hepatocytes to first induce senescence and then later, develop into cancer (Mudbhary *et al*., 2014). We identified DSB, ATM, and Tp53 activation as a mechanism of senescence caused by hUHRF1 overexpression, potentially mediated by DNA hypomethylation of TEs and their activation of LTR transposons. Moreover, we show that this response occurs within hours of UHRF1 overexpression, indicating that even a short-term disruption in UHRF1 expression can induce cancer-causing changes. We identify multiple senescent phenotypes induced by hUHRF1 overexpression, and postulate that the widespread DNA methylation changes caused by deregulated UHRF1 levels can disrupt multiple pathways. We further show that these different senescent cell phenotypes evolve, so that at early stages senescence is sustained even if *atm* and *tp53* are removed. At later stages, a population of cells emerges with distinct gene expression profiles and with reduced UHRF1 levels; these can receive proliferative signals and, if *tp53* is lost, can reenter the cell cycle, indicating senescence escape. Another population of cells with the highest levels of UHRF1 cannot proliferate, but signal to neighboring cells and support the microenvironment in this precancer state. Importantly, our data indicate that senolytic therapy may remove only one subset of senescent cells, allowing those with escapable senescence to proliferate.

Senescence occurs when cells in a senescent state that retains the ability to reactivate proliferative genes lose key tumor suppressive signals (Evangelou *et al*, 2023). Our finding that hepatocyte proliferation increases during later stages of the senescence-to-cancer transition suggests senescence escape. This is reminiscent of a mouse model of HCC, whereby Ras overexpression induced senescence, and the Ras expressing cells were rapidly cleared by the immune system. However, when immune cells were depleted, Ras-expressing cells persisted and progressed to form liver cancer (Kang *et al*., 2011). A more recent study demonstrated that this is attributed to the level of Ras, which can act as a potent oncogene in HCC only when expressed at low levels, because hepatocytes with high Ras expression acquire an immunogenic senescence phenotype and are rapidly cleared. In contrast, cells with lower Ras expression have a senescence phenotype that is immune-evasive and escapable, leading to HCC (Chan *et al*., 2024). This phenomenon parallels our observations with UHRF1. High UHRF1 levels induce senescence, functioning as an effective tumor-suppressive barrier; these cells remain non-proliferative even when the ATM-Tp53 pathway is inactivated. However, a subset of senescent cells reduces UHRF1 expression through an unknown mechanism while retaining epigenetic alterations and DNA damage. Upon *Tp53* loss, these cells can re-enter the cell cycle, suggesting a potential route for malignant transformation. Thus, UHRF1-induced senescence has a dual role, initially restraining tumorigenesis but later creating conditions that may promote cancer initiation.

How the epigenome is repatterned in cancer is not known. One possibility is that DNA methylation changes occur during oncogene induced senescence as a byproduct of the process (Cruickshanks *et al*., 2013; Hlady *et al*., 2017; Xie *et al*, 2018). This could be due to deregulation of epigenetic writers and readers, as loss of UHRF1 has been shown to be sufficient to induce senescence (Chen *et al*., 2025; Jung *et al*., 2017). It is interesting that even *in vitro*, UHRF1 loss in cancer cells induces diverse senescence phenotypes over time (Chen *et al*., 2025). In cancer cells, UHRF1 depletion causes senescence likely driven by DNA hypomethylation as well as potentially due to DNA methylation–independent functions of UHRF1. Future studies to determine whether the senescent phenotypes arising from UHRF1 overexpression and loss are overlapping or distinct will be an important focus. *In vivo*, we speculate that even transient deregulation of UHRF1 levels could change DNA methylation patterns during DNA replication, either by titration or degradation of Dnmt1 (Mudbhary *et al*., 2014) and by recruitment of Dnmt3a or by repelling Tet2-mediated demethylation to promote hypermethylation (Yamaguchi *et al*., 2024). In such cases, either gain or loss of UHRF1 could induce senescence, which could then result in clearing the resulting cells with epigenetic damage. In the case where these cells are not cleared, the possibility of senescence escape or persistence and signaling to neighboring cells as we observed here could be tumor promoting.

An unresolved question is the source of UHRF1 heterogeneity, despite its expression being driven by a strong promoter. This may result from epigenetic silencing of the locus, the selective expansion of rare cells that have lost the transgene, or a shift toward a more stem-like identity with reduced hepatocyte differentiation and thus the downregulation of UHRF1 expression by the fabp10a promoter. While the mechanism remains unclear, it provides a useful system since the heterogeneity of UHRF1 mirrors the variable expression observed in human HCC (Mudbhary *et al*., 2014).

The work presented here raises considerations for understanding the nature of senescence and targeting therapies. First, the finding that dramatic changes to the transcriptome, including TE activation, occurs within 1 day of UHRF1 overexpression in this model, accompanied by DNA damage indicate that even a modest deregulation of UHRF1 can have dramatic, cancer-causing effects. In these cases, targeting early senescence may be advantageous in preventing cancer. However, approaches to selectively reinforce tumor-suppressive senescence while eliminating pro-tumorigenic senescent cells using senescent promoting (senogenic) and senescent eliminating (senolytic) drugs (Colucci *et al*., 2025) have not been translated to a standard of care for any type of cancer. While the hope that such therapies can harness the tumor suppressive features, the pleiotropic and context-dependent nature of senescent cell phenotypes means that both tumor suppressing and tumor promoting senescent cell states can be induced by the same oncogene and thus exist in the same tumor. Therefore, applying these drugs to activate the anti-cancer senescent phenotypes and restrict those that promote the tumor ecosystem relies on understanding how senescent cell identities are generated and can be manipulated.

### Material and Methods Zebrafish Husbandry

Zebrafish (*Danio rerio*) care was conducted according to New York University Abu Dhabi (NYUAD) for Animal Care and Use Committee (IACUC) Committee (protocol number 22-0003A2). Adult fish were raised on a 14:10 h light: dark cycle at 28 °C and fed 2 times a day with artemia and once with Zeigler Zebrafish Diet. Larvae after 5 dpf were fed two times daily with paramecia until 3-4 weeks when they were started on artemia. *Tg(fabp10a:hsa.UHRF1-EGFP^mss1^)^high^*(Mudbhary *et al*., 2014) called *hUHRF1*, Tg(*fabp10a:CAAX-EGFP)* (Nussbaum *et al*, 2013), Tg(*fabp10a:nls-mCherry)* (Mudbhary *et al*., 2014), Tg(*mpeg1.1:dsRed2)* (see below), and *Tg(lyzC:dsRed)(Hall et al, 2007)* were used to identify macrophages and neutrophils, respectively. *tp53^−/−^* (Berghmans *et al*., 2005) and *atm^−/−^*(Ajouaou *et al*., 2023) mutant zebrafish were raised as incross and genotyped by PCR to identify homozygous *tp53^−/−^* or genotyped by Sanger sequencing to identify homozygous *atm^−/−^* mutants. *Tg(fabp10a:hsa.UHRF1-EGFP^mss1^);atm^−/−^*or *Tg(fabp10a:hsa.UHRF1-EGFP^mss1^);tp53^−/−^* and were outcrossed to *atm^−/−^, tp53^−/−^*, or WT fish respectively so that in all experiments, the *hUHRF1* allele was heterozygous.

### Generation of zebrafish line

*Tg(mpeg1.1:dsRed2)* fish expressing dsRed2 in macrophages was generated by PCR amplification of dsRed2 (Yoshinari *et al*, 2012). BamH1 (New England Biolabs) and MfeI (New England Biolabs) restriction enzyme sites were inserted with PCR (Q5 Taq, New England Biolabs) in the forward and reverse primer respectively. A vector containing tol2 sites and mpeg1.1:Dendra2 cassettes (Addgene: #51462) was digested with the same restriction enzymes. After ligation and transformation in DH5alpha (Thermo Fisher Scientific), colonies were screened by PCR for the presence of dsRed2. Positive colonies were sequenced to avoid the presence of point mutations. One clone was selected and amplified by GenElute Plasmid Miniprep kit (Sigma Aldrich) and further purified with MinElute PCR Purification Kit (Sigma Aldrich). 1 nl of 40 ng/ml of plasmid was injected with 80 ng/ml ng of Tol2 transposase mRNA into 1-2 cell stage embryos. Once embryos reached sexual maturity, they were crossed to WT fish to generate F1 and selected for bright expression of dsRed2 in macrophages.

### Drug treatment of zebrafish larvae

All treatments were performed on at least 10 larvae from control and *hUHRF1* samples in 6 well plates containing 10 ml of embryo medium and incubated at 28 °C with 14:10 dark/light cycle according to standard protocols (Ramdas Nair *et al*, 2021). VE-821 (ab219506, Abcam) was dissolved in DMSO (Sigma) at 20 mM stock solution stored at -20 °C. Larvae were treated with 10 μM VE-821 or 0.05 % DMSO from 72 to 96 or 120 hpf when larvae were collected for EdU or SA-β-gal staining, respectively. Dasatinib (Sigma) was resuspended in DMSO to prepare 20 mM stock and Navitoclax (Sigma) was resuspended in DMSO to prepare a 10 mM and stored at -20 °C in aliquots. Larvae were treated from 4 to 7 dpf with a final concentration of 2.5 μM Dasatinib or with 25 μM Navitoclax in 1 % DMSO. DMSO 1% was used as vehicle. At 7 dpf larvae were collected for EdU or SA-b-galactosidase analysis.

### EdU incorporation

*hUHRF1* and control larvae at the indicated time points were incubated in 250 ml of embryo medium containing 10% DMSO and 0.5 mM EdU (Thermo Fisher Scientific, resuspended in embryo water) for 20 minutes on ice. After 20 minutes, 50 ml of embryo medium was added to the larvae and then incubated at 28 °C for 30 minutes. For 7 and 10 dpf larvae, 0.22 μm filtered system water was used instead of embryo medium for the EdU incorporation. After EdU, larvae were fixed with 4% paraformaldehyde overnight at 4 °C, washed once with PBS and gradually dehydrated in 100% methanol and stored at 4 °C overnight. EdU pulsed larvae were stained by Click-IT (Thermo Fisher Scientific) as described (Madakashira *et al*, 2023). Briefly, larvae were gradually rehydrated in 100 % PBS and incubated twice with PBS containing 8 mM CuSO_4_ (Sigma Aldrich), 4 nM Sodium Azide (A555 or A647, Thermo Fisher Scientific) and 50 mM Ascorbic Acid (Sigma Aldrich) for 30 minutes at room temperature. Livers were dissected, stained with Hoechst (1:2000, Thermo Fisher Scientific) and GFP booster (1:1000, Chromotek) for 1 hour at room temperature and washed 3 times in PBS before imaging. A z-stack through each liver (z = 0.33mm) was acquired at Leica Stellaris 8 Confocal Microscope using the Lightning function. Images were quantified by Imaris using the Spot function to count Hoechst, EGFP and EdU positive cells. To determine the level of EGFP in the EdU positive cells, the intensity of EGFP in EdU positive cells that were identified by the Spot function was measured in pixels. For each liver, the pixel intensity of EGFP signal for the 3 brightest EGFP positive cells, regardless of EdU status, was measured and used to normalize the EGFP signal to set the top level. For each time point and condition, at least 3 independent biological replicates were used.

### Immunofluorescence

*hUHRF1* and control larvae from indicated time points were collected and fixed with 4% paraformaldehyde at 4 °C overnight and processed as described (Madakashira *et al*., 2023). Briefly, larvae were washed twice with PBS, livers were manually dissected and incubated with PBS containing 1 % Triton-X (Sigma Aldrich) and 20 mg/ml of Proteinase K (Machery-Nagel) for 10 minutes at room temperature then washed with PBS and incubated for 1 hour at room temperature with PBS containing 2% bovine serum albumin (Sigma Aldrich) followed by incubation at 4 °C overnight with primary antibody in blocking solution. After primary antibody staining, livers were washed 3 times with PBS containing 0.1% Tween-20 and 2 washes with PBS, followed by incubation with secondary antibody solution (blocking solution with 1:300 secondary antibody and 1:2000 Hoechst, Thermo Fisher Scientific) for 1 hour at room temperature. After washing with PBS, livers were mounted with Vectashield (Vector) and imaged on a Leica Stellaris 8 Confocal Microscope and quantified with LasX software. Primary antibodies were γH2AX (gtx127342, Genetex), p-RPA (ab211877, AbCam) at 1:100. Secondary antibodies were anti-rabbit Alexa 555 (Thermo Fisher Scientific), anti-mouse Alexa 546 (Thermo Fisher Scientific), anti-rabbit Alexa 647 (Thermo Fisher Scientific), anti-mouse Alexa 647 (Thermo Fisher Scientific). At least 3 livers were imaged for each experiment in at least 2 clutches.

### Analysis of fluorescence heterogeneity

*hUHRF1* and nls-mCherry lines were crossed and *hUHRF1;nls-mCherry* or *nls-mCherry* larvae were collected at indicated time points and fixed in 4 % paraformaldehyde at 4 °C overnight and then transferred in PBS. Livers were manually dissected, permeabilized with PBS containing 1 % Triton-X (Sigma Aldrich) and 20 μg/ul Proteinase K (Machery-Nagel) for 10 minutes at room temperature and stained with 1:2000 Hoechst (Thermo Fisher Scientific) in PBS for 1 hour at room temperature. Livers were washed 3 times with PBS and were mounted on a glass slide with Vectashield (Vector). Images were acquired with Leica Stellaris 8 Confocal Microscope. For each liver, the laser power was set to have 1-2 saturated cells for each channel. Image analysis was performed with Fiji by creating a mask on the Hoechst channel and then quantifying the intensity of mCherry and EGFP for each cell present in the mask. For each image and for each channel, intensity levels of each cell were normalized based on the cell with the highest intensity in that image and relative intensity was plotted for visualization in GraphPad Prism 8.

### SA-β-gal staining

SA-β-gal staining was performed using a kit (Cell Signaling Technologies). Briefly, *hUHRF1* or control larvae at 5 dpf or 7 dpf were collected and fixed with fixing solution for 15 minutes at room temperature followed by 3 washes with PBS pH 7.4. Staining was performed overnight at 37 °C with staining solution with optimized pH at 6.2. Larvae were washed in PBS and mounted in 3 % methylcellulose for imaging at stereoscope. SA-β-gal staining in the posterior gut was considered as positive control for staining and larvae were scored blindly for the presence of the staining in the liver and categorical assigned as positive or negative.

### RNA extraction, library preparation and bulk RNAseq analysis

Pools of 20-30 manually dissected livers from 5 independent clutches of *hUHRF1* and control larvae at 80 hpf or 10-15 livers at 120 hpf were used for RNA isolation. Livers were collected directly in 500 ml of Trizol (Thermo Fisher Scientific). RNA was extracted by following manufacturer’s instruction with some modification. After chloroform, RNA was precipitated in isopropanol (Sigma Aldrich) with 10 μg of Glycoblue (Thermo Fisher Scientific) overnight at – 20 °C. After precipitation, RNA was centrifuged at 4 °C for 1 hour, washed in 70 % ethanol and resuspended in 20 μl of DNAse/RNAse free water. Possible genomic DNA contamination was removed with RapidOut DNA removal Kit (Thermo Fisher Scientific) for 30 minutes at 37 °C.

After quality control with Bioanalyzer (Agilent), 100 ng of RNA was used for library preparation with ILMN Strnd Total RNA Lig w/RBZ+ (Illumina). Libraries were sequenced on NovaSeq 6000 Illumina platform. Quality of the sequences was assessed by using MultiQC v1.0 (https://multiqc.info). After adaptor removal and trimming, reads were aligned to the *D. rerio* GRCz10 reference genome, with manual insertion of UHRF1 and EGFP sequence, using HISAT2 with default parameters (Kim *et al*, 2019), mapped and counted with HTSeq (Anders *et al*, 2015). Differential gene expression was calculated using a generalized linear model implemented in DESeq2 in Bioconductor (Love *et al*, 2014) to test differential gene expression between *hUHRF1* transgenic livers and WT sibling controls or, in the case of *tp53^-/-^* mutants, normalized to *tp53^-/-^*samples. Adjusted p-value with a false discovery rate of <0.05 was used to determine DEGs.

Quantification of repetitive elements was assessed by SQuIRE workflow (Yang *et al*, 2019) to quantify expression at the subfamily level and properly assign multi-mappers sequences (due to the nature of repetitive elements) using expectation-maximization algorithm as described (Madakashira *et al*., 2023). In brief, trimmed fastq were aligned to D. rerio GRCz10 reference genome using STAR. REs annotation from RepeatMasker was used for alignment with unique mapping reads aligned to a single locus and marking multi-mapped reads to multiple locations and further refined with EM algorithm with “auto” parameter. Total counts (unique and multi-mapped) are aggregated by subfamily of REs and the output count table from SQuIRE has been analyzed by DEseq2 with standard parameters to calculate differential expression of REs between condition.

For gene ontology analysis, DEG (p-value adjusted < 0.05) at 80 hpf and 120 hpf were used to perform Ingenuity Pathways Analysis (Qiagen) and top induced pathways at 80 hpf were plotted using Graph Prism 8. Heatmaps were generated with Pheatmap on Log2FC of *hUHRF1* compared to controls and plotted in R.

### DNA extraction, RRBS and DNA methylation analysis

Genomic DNA was extracted from 80-100 livers dissected from 3 independent clutches of *hUHRF1* or Tg(fabp10a:CAAX-EGFP) larvae as controls using a DNA extraction buffer (10 mM Tris-HCl pH9, 10 mM EDTA, 200 mM NaCl, 0.5% DSD, 200 μg/ml proteinase K). DNA was resuspended in water and quantified by Qubit dsDNA High Sensitivity kit. RRBS was performed on ∼80 ng of genomic DNA and digested with 200 U of MspI (New England Biolabs) and used for library preparation with minor modifications on previously published protocol (Magnani *et al*., 2021) (Garrett-Bakelman *et al*, 2015). To avoid loss of gDNA, after MspI digestion, end repair, and A-tailing, the reactions were stopped by heat inactivation. The adaptors used for multiplexing were purchased separately (Next Multiplex Methylated Adaptors-New England Biolabs). Libraries were size-selected for fragments from 175 bp to 670 bp by dual-step purification with Ampure XP Magnetic Beads (Beckman Coulter). Bisulfite conversion was performed with Lightning Methylation Kit (ZYMO Research) following the manufacturer’s instructions. Libraries were amplified using KAPA HiFi HotStart Uracil+ Taq polymerase (Roche) and purified with Ampure XP Magnetic Beads (Beckman Coulter) before sequencing. Libraries were sequenced as 150 bp paired end reads on the Illumina Nextseq 600 platform. Quality control of the RRBS sequencing data was assessed using FASTQC and Trimmomatic and aligned to the reference genome GRCz10 (Bolger *et al*, 2014).

RRBS data was analyzed for CpG methylation levels using the R package ‘methylKit’ (Akalin *et al*, 2012). CpGs covered at least 10 times in one biological replicate were included in the analysis. CpGs with a q-value indicating a false discovery rate of <0.01 were considered differentially methylated, and they were segregated between those that had a methylation change of greater or less than 25% in hUHRF1 samples compared to controls. For plotting and statistical analysis, R package ‘pheatmap’ and GraphPad Prism software were used.

### Preparation of single cell suspension and scRNAseq

150-800 livers at 5 dpf from Controls and *hUHRF1* were manually dissected and put in ice-cold PBS containing 2 % FBS (Gibco) and 1 % penicillin/streptomycin (Gibco). After dissection, livers were briefly centrifuged, resuspended in 1 ml of Trypsin/EDTA 0.05% (Sigma Aldrich) and incubated at 28 °C for 12-20 minutes. Every 2 minutes livers were pipetted 10 times to facilitate cell dissociation until all the cells resulted in a homogenous suspension. 200 μl of FBS (Gibco) was added to inactivate Trypsin and cells were centrifugated at 4 °C for 7 minutes at 500 g. After centrifugation cells were washed 3 times with PBS containing 0.4 % BSA (Sigma Aldrich). Cells were resuspended in 50-100 μl of PBS containing 0.4% BSA and counted with a Burker Chamber. Cell suspension at a concentration of 600-700 cells/μl was used to load Chip for 10X Genomics acquisition. All steps of single-cell sequencing were performed following the Chromium Next GEM Single Cell 3ʹ Reagent Kits v3.1 (Dual Index).

### Single cell RNAseq analysis

scRNAseq data were processed and filtered using Seurat v5.0.1 (Hao *et al*, 2024). Initial quality control and filtering of scRNAseq data were performed to ensure the retention of high-quality cells. In that line, cells with fewer than 200 detected features and nCount_RNA or mitochondrial gene content greater than 20% were excluded from downstream analysis. Data filtering resulted in 12,202 cells for Controls and 10,919 cells for *hUHRF1*. Highly variable features were identified using the "vst" method with the top 2,000 features selected. Data integration was performed with Seurat package using the FindIntegrationAnchors function with canonical correlation analysis (CCA)(Stuart *et al*, 2019). Dimensionality reduction and clustering of scRNAseq data were performed using principal component analysis (PCA), retaining the top 19 principal components. Uniform Manifold Approximation and Projection (UMAP) dimensionality reduction was applied using these components for visualization. Neighbor identification was conducted with FindNeighbors (k = 20), followed by clustering using FindClusters with a resolution of 0.3. DEGs in each cell cluster were identified using FindMarkers function, and statistically significant markers between compared groups were selected based on an adjusted p-value threshold (p-value < 0.05) were used to perform gene ontology. All dimensionality reduction and DEG analyses were conducted using the Seurat package.

To define differences between population, ClusterProfiler package was used in R. Significantly enriched markers (p-value adjusted < 0.05 & Log2FC > 0.5) were used to perform Gene Ontology. The 3 most significant GO terms were plotted by dotplot function in ClusterProfiler.

To define the differences between *hUHRF1* and controls, differential gene expression was performed on *hUHRF1* compared controls for each population. DEGs (p-value adjusted < 0.05 & Log2FC of 1.5) in each hepatocyte population were used to perform KEEG pathway analysis using ClusterProfiler package in R. The 5 most significant KEEG terms were plotted using dotplot function in ClusterProfiler. To identify high, mid and low expressing cells, hepatocyte population in *hUHRF1* and controls were selected and quantiles of EGFP were defined. Cut off of EGFP separating the 10 % most expressing cells (HIGH) and 10 % least expressing cells (LOW) were selected and MID cells were defining as cells expressing the middle 11-80^th^ percentile of EGFP levels. DEGs were identified using FindMarkers function on High, Low and Mid cells down sampled (seed 123) to match the number of cells in High and Low (1416 cells) and separated in controls and *hUHRF1*.

### Cell-cell communication analysis

Cell-cell communication was inferred at single-cell resolution using the CellChat R package (v2.1.2) following the standard workflow (Jin *et al*, 2021). Ligand-receptor interactions from secreted signaling were mapped using the zebrafish-specific CellChatDB.zebrafish database. Communication probabilities were estimated with *computeCommunProb* and interaction pairs represented by fewer than ten cells were excluded. Pathway-level signaling probabilities were obtained with *computeCommunProbPathway* function and summarized using *aggregateNet*. Interaction numbers and strengths were visualized as chord diagrams using *netVisual_chord_cell*.

Outgoing and incoming signaling roles were quantified with *netAnalysis_computeCentrality* and displayed using *netAnalysis_signalingRole_heatmap*.

## Statistical analysis

Statistical analysis was performed in GraphPad Prism 8. At least 3 biological replicates were used for each experiment. Methods to evaluate the statistical significance include two-tailed Student’s *t*-test with adjustment for multiple comparisons, t-test, or Chi-square for categorical variables. Tests used are indicated in figure legend. All the plots were generated in GraphPad Prism 8 and RStudio 3.3.1.

## Acknowledgements

This work was supported by NYUAD Faculty Research Fund (AD188), NIH (5R01DK080789-11) to KCS and Tamkeen under the NYU Abu Dhabi Research Institute Award to the NYUAD Center for Genomics and Systems Biology (ADHPG-CGSB) and the Al Jalila Foundation (to KCS). All imaging was carried out in the NYUAD Core Technology Platform Imaging facility with expert support by Rashid Razgui. Bulk RNAseq and scRNAseq libraries were performed by NYUAD Core Technology Platform Imaging with expert help of Marc Arnoux and Mehar Sultan. Bioinformatics assistance was provided by the NYUAD Bioinformatics Core at New York University Abu Dhabi. We would like to thank Nizar Drou, Muhammad Arshad and Giuseppe-Antonio Saldi for expert bioinformatics support. We thank all the member of the Sadler lab, in particular Shashi Ranjan for contribution to animal maintenance and experiments and Momal Taimoor for editing the manuscript.

## Author contributions

EM and KCS conceptualized the project; EM, CC, FM, BM, IMB and TR developed the methodology and carried out the experiments and analysis; EM, FM, CC and TR created the visualization, KCS provided resources, supervision, project administration, wrote the first draft and EM and KCS reviewed and edited the manuscript. All the authors read and approved the final version of the manuscript.

## Data availability

Raw data (fastq files) and processed files (raw counts) for bulk RNAseq have been deposited in GEO as GEO: GSE302812. RRBS data has been deposited in GEO in GEO: GSE173792. Raw data (fastq files) and processed files (Cell ranger output) for single cell-RNAseq have been deposited in GEO as GSE303361.

This paper does not report original code.

---

**Table S1.** DEGs in bulk RNAseq at 80 hpf and 120 hpf in hUHRF1 compared to wild type siblings.

**Table S2.** CpG methylation changes induced by hUHRF1 overexpression.

**Table S3.** DEGs in bulk RNAseq at 120 hpf in hUHRF1 compared to wild type siblings, in *tp53^-/-^;hUHRF1* compared to *tp53^-/-^* and *in tp53^-/-^* compared to wild type livers.

**Table S4.** Markers of each population identified in scRNAseq merging hUHRF1 and controls.

