## Supplemental Information for "UHRF1 Overexpression Generates Distinct Senescent States with Different Tp53 Dependencies"

### Supplemental Figures

#### Figure S1

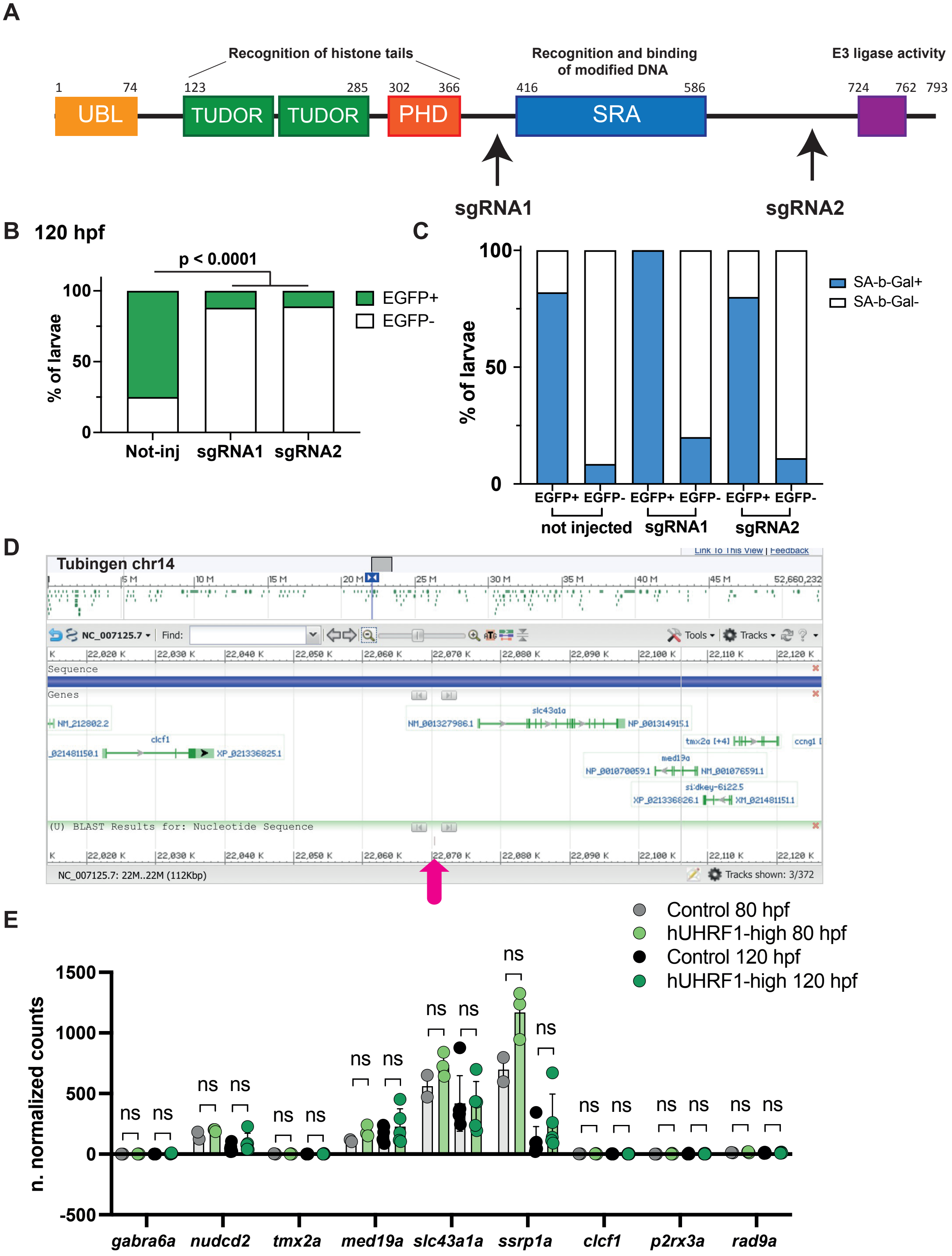

**Supplemental Figure S1. Senescence in the hUHRF1 is independent of the transgene insertion site. A.** Schematic representation of hUHRF1, its domains and the region targeted by sgRNAs. **B.** Percent of larvae with GFP expression as a marker of hUHRF1-EGFP mutation. **C.** SA-β-gal staining of non-injected and injected larvae. **D.** Chromosomal view of hUHRF1 insertion. **E.** Expression (by bulk RNAseq) at 80 and 120 hpf of genes flanking the insertion site.

**Figure S2**

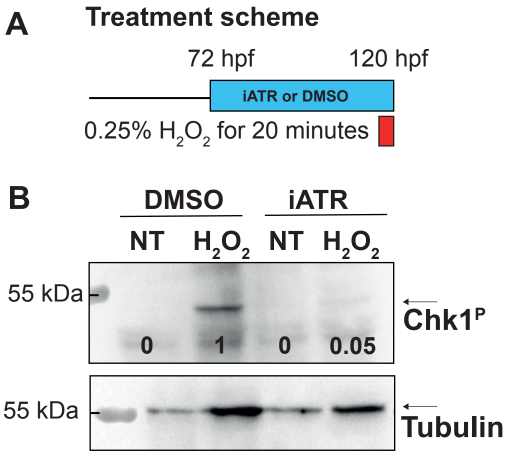


**Supplemental Figure S2. Validation of VE-821 as Atr inhibitor block. A.** Treatment scheme of ATR inhibitor performed in *hUHRF1* and Control larvae. To determine the efficacy, $H_2O_2$ was used as DNA damage inducer. **B.** Western blot of Control and $H_2O_2$ treated larvae in presence and absence of the Atr inhibitor VE-821.

#### Figure S3

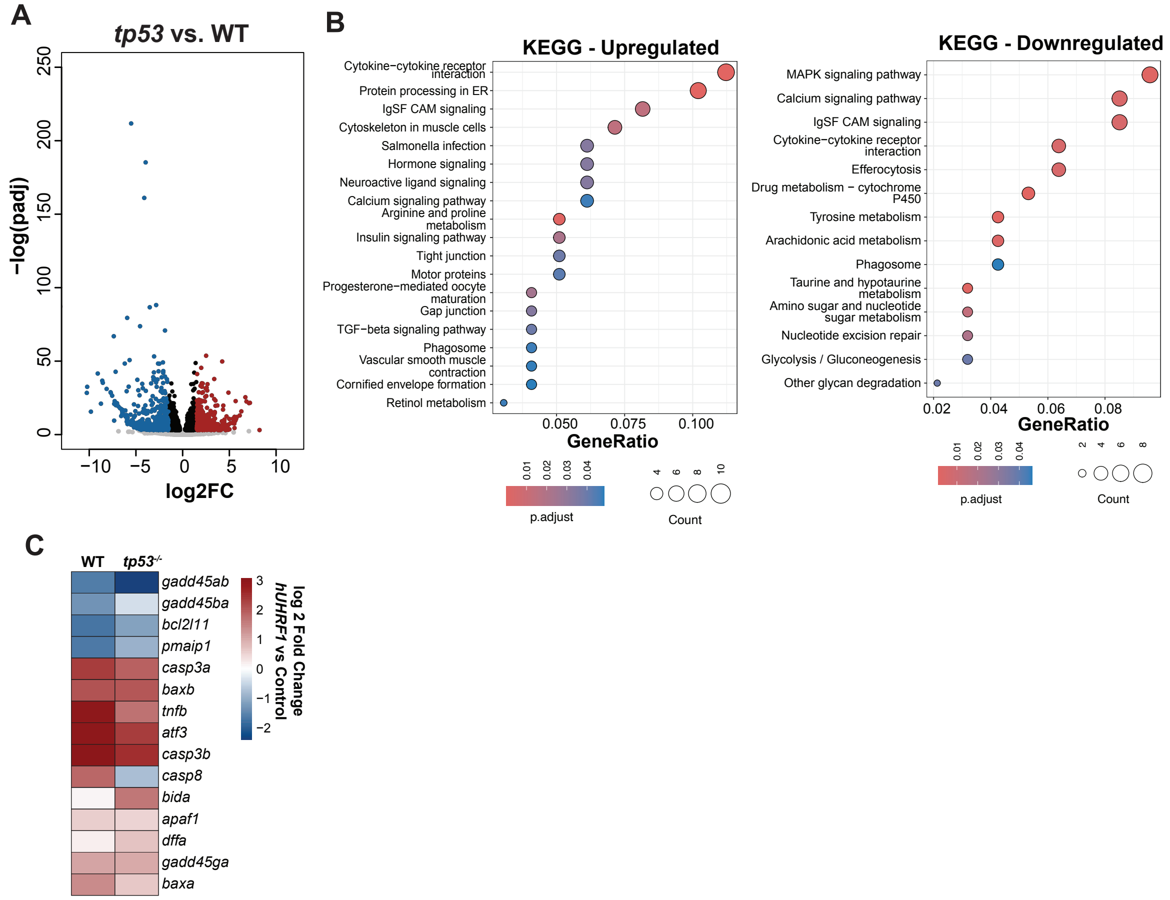


**Supplemental Figure S3. Bulk RNAseq of *tp53*$^{-/-}$ compared to wild type. A.** Volcano plot of DEGs between *tp53*$^{-/-}$ compared to wild type. **B.** KEEG pathway analysis of upregulated genes. **C.** KEEG pathway analysis of downregulated genes. **D.** Heatmap of Log2FC of genes involved in apoptosis in *hUHRF1* compared wild type and *tp53*$^{-/-}$; *hUHRF1* compared to *tp53*$^{-/-}$.

**Figure S4**

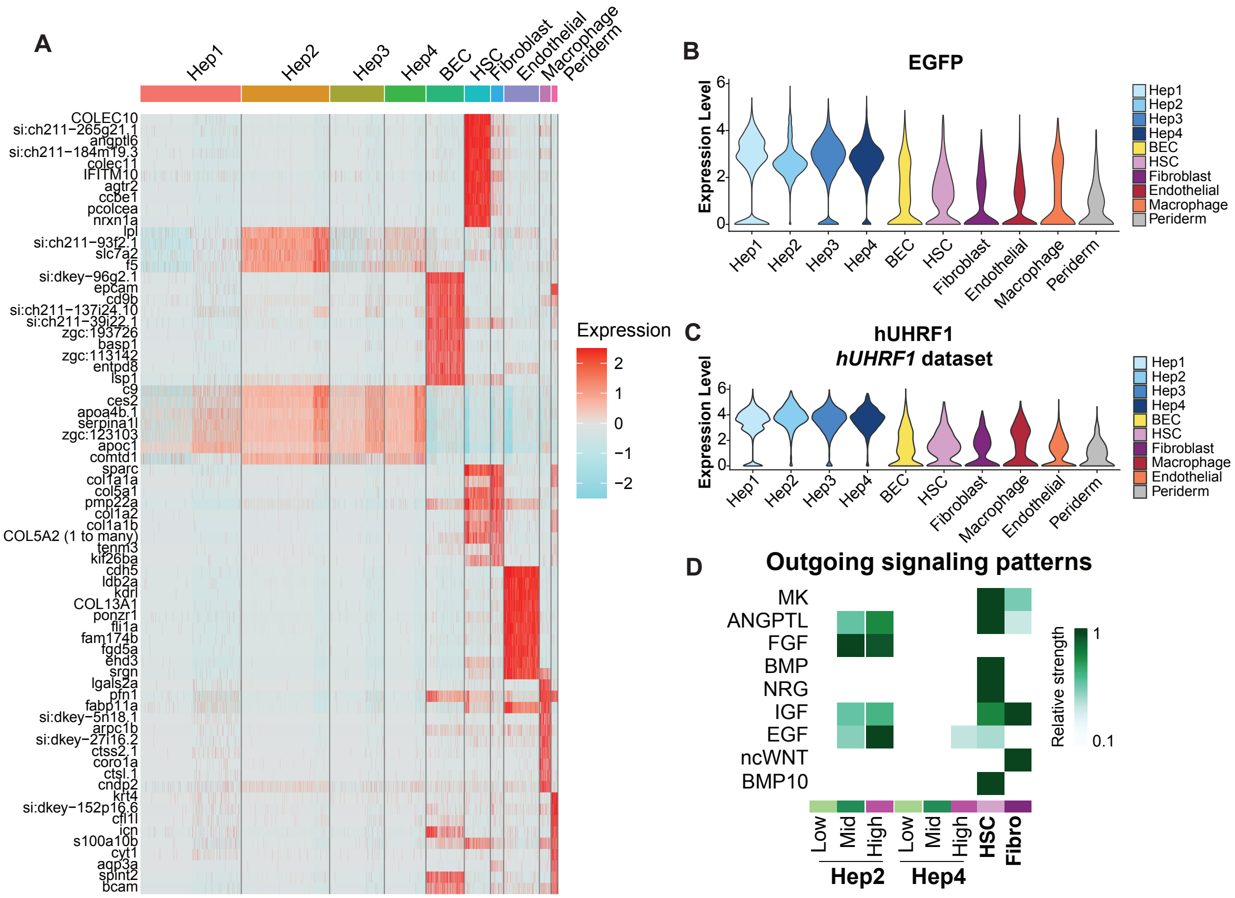


**Supplemental Figure S4. Single-cell RNAseq analysis**. **A.** Heatmap of key identity markers in different cell populations. **B.** Violin plot of EGFP levels across populations. **C.** Violin plot of hUHRF1 levels across populations. **D.** Heatmap of outgoing signaling pathways in Hep population divided in high, mid, and low, mesenchymal cells and macrophages in *hUHRF1* samples.

**Figure S5**

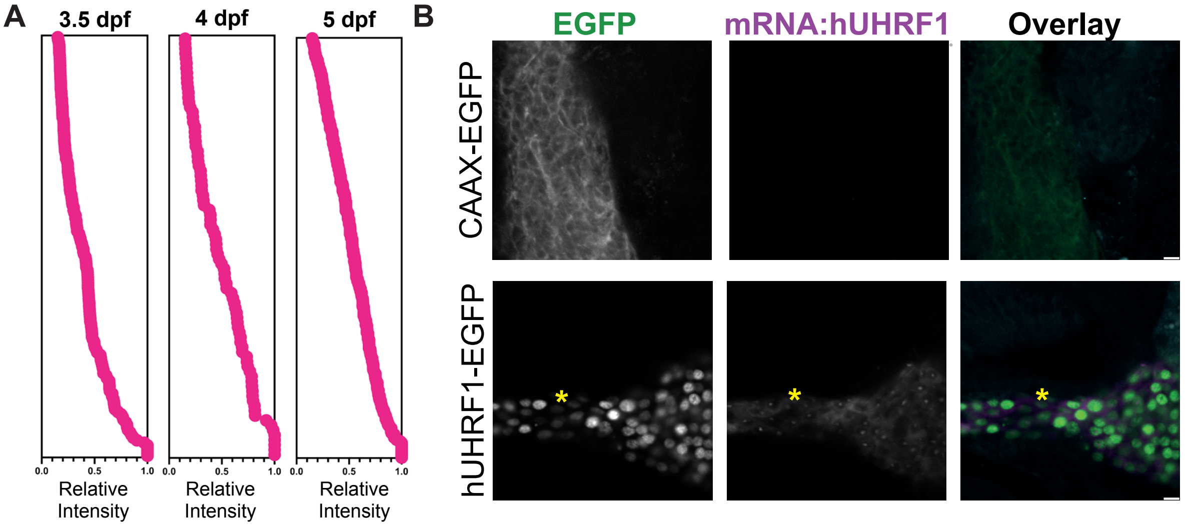


**Supplemental Figure S5. *hUHRF1* expression is heterogenous in hepatocytes**.

**A.** Quantification of nls-mCherry signal in controls. **B.** In situ hybridization of hUHRF1 shows that cells with low/no GFP also have low/no hUHRF1 mRNA.

#### Supplementary Materials and Methods

##### Crispr/Cas9 deletion of human UHRF1

sgRNA targeting the human UHRF1 gene was produced by sgRNA IVT kit (Takara Bio), the resulting RNA was isolated using Trizol (Invitrogen) and was quantified by Qubit. The sgRNA was diluted to 50 ng/µL, mixed with an equal volume of previously diluted nls-Cas9 protein (IDT; 0.5 µl of nl-Cas9 added with 9.5 µL of 20 mM HEPES; 150 mM KCl, pH 7.5) and incubated at 37 °C for 5 min. We then injected 1 nl into 1–2 cell stage embryos from an incross of *hUHRF1* which were then raised to 120 hpf. At 120 hpf embryos were screened for the presence of EGFP in the liver in both not injected and injected Controls. 75 % positive is the expected percentage of EGFP in not-injected controls while the injected, if crispants, should show a decreased percentage. EGFP positive and negative from each group were used to perform SA-ß-galactosidase staining.

##### Mapping fabp10a:hUHRF1-EGFP integration

gDNA from hUHRF1-EGFP positive zebrafish was extracted as described. 5 µg of gDNA was digested with AluI (New England Biolabs) at 37 °C for 3 hours. Digested DNA was purified with PCR purification kit (Sigma Aldrich) then ligated with T4 DNA ligase (New England Biolabs) at 16 °C overnight. Ligated products were PCR amplified with PCR followed by nested PCR by targeting the 5' and 3' region of the cassette directed outside the cassette. Water was used as PCR control and WT gDNA was processed in parallel as negative control. PCR product was run on a 1 % agarose gel and interested band was gel purified with Gel Elute kit (Sigma Aldrich). Purified products were sequenced and mapped to zebrafish genome by nBLAST.

##### In situ HCR (Hybridization Chain Reaction)

hUHRF1 and Controls (CAAX-EGFP) from 5 dpf larvae were fixed in 4 % paraformaldehyde overnight at 4 °C, washed 3 times with PBS and gradually de-hydrated in 100 % methanol. After gradual rehydration in 100 % PBS in situ HCR was performed as per the manufacturer's instructions without modifications (Molecular Instruments). After washing, whole larvae were stained for 1 hour at room temperature with Hoechst (1:2000 in PBS, Thermo Fisher Scientific) and washed 3 times with PBS followed by mounting in 3 & methylcellulose. Imaging was performed at confocal microscope Stellaris.
